# Layer-specific sensory processing impairment in the primary somatosensory cortex after motor cortex infarction

**DOI:** 10.1101/778167

**Authors:** Atsushi Fukui, Hironobu Osaki, Yoshifumi Ueta, Yoshihiro Muragaki, Takakazu Kawamata, Mariko Miyata

## Abstract

Primary motor cortex (M1) infarction occasionally causes sensory impairment. Because sensory signal plays an important role in motor control, sensory impairment compromises recovery and rehabilitation from motor disability. Despite the importance of sensory-motor integration for rehabilitation after M1 infarction, the neural mechanism of the sensory impairment is poorly understood. We show that the sensory processing in the primary somatosensory cortex (S1) was impaired in the acute phase of M1 infarction and recovered in a layer-specific manner in the subacute phase. This layer dependent recovery process and the anatomical connection pattern from M1 to S1 suggested the functional connectivity from M1 to S1 plays a key role in the impairment of sensory processing in S1. The simulation study demonstrated that the loss of inhibition from M1 to S1 in the acute phase of M1 infarction could cause the sensory processing impairment in S1, and the complementation of inhibition could recover the temporal coding. Taken together, we revealed how focal stroke of M1 alters cortical network activity of sensory processing, in which inhibitory input from M1 to S1 may be involved.

## Introduction

Sensory information strongly influences motor coordination.^1^ Consistently, it is also essential for the restoration of motor performance after stroke and is frequently used for effective neurorehabilitation.^2,3^ However, primary motor cortex (M1) infarction leads to not only motor dysfunction, but also somatosensory impairment in the acute phase after M1 infarction in primates.^4,5^ Because rehabilitative therapies are most beneficial when initiated in the acute phase of stroke,^6^ we need to understand how sensory processing is modified after M1 infarction, especially shortly after injury. Despite many studies on the neural mechanism, including the reorganisation and recovery process of motor dysfunction after M1 infarction,^7,8^ the fundamental spatiotemporal dynamics of sensory processing in the primary sensory cortex (S1) remain unknown.

To address this issue, we used the photothrombotic method on mouse vibrissa M1 (vM1) as a model of focal ischemic stroke and investigated the effect on somatosensory processing in the mouse vibrissa S1 (vS1). vS1 is spatially separated from vM1^9^ and ideal to observe a pure M1 infarction effect. We examined the response reliability to whisker stimulation from each vS1 layer in the acute and subacute phases after the vM1 infarction and found impairment of sensory processing in all layers in the acute phase, but recovery in the subacute phase except in the deep vS1 layer, where vM1 innervates densely. We also conducted a simulation study to predict the circuit mechanisms for the sensory processing impairment in vS1.

## ResultA photothrombotic infarction model in vM1

We made the vM1 infarctions by using the photothrombotic method and recorded MUA from vS1 at POD3 and POD14 (Fig. 1A). The sizes of the infarction areas centred at the vM1 were stable on each experimental day, but were significantly smaller at POD14 than at POD3. The means of the largest normalized areas were 1.52±0.26 mm^2^ at POD3 (N = 5) and 0.74±0.08 mm^2^ at POD14 (N = 4, P = 0.02, two sample t-test, Fig. 1B). This reduction was also observed in other studies using photothrombotic infarction.^10,11^

**Figure 1.**
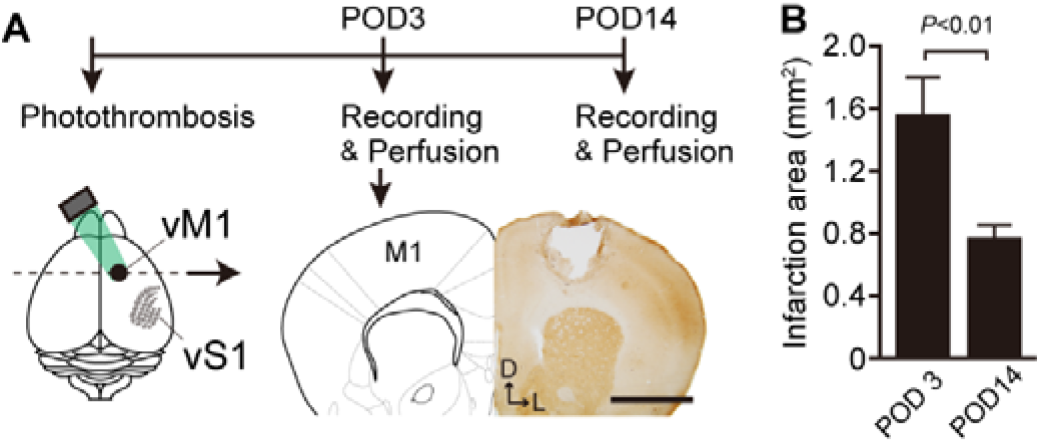
Experimental timeline of the vM1 photothrombotic infarction model **A**, Mouse vibrissa primary motor cortex (vM1) infarction model made by the local irradiation of green light. Electrophysiological recordings from vibrissa primary somatosensory cortex (vS1) were performed at postoperative day 3 (POD3) and POD14. *Right*, The infarction site was identified by cytochrome oxidase staining (1.4 mm anterior to the bregma, POD3). Scale bar, 1 mm; D, dorsal; L, lateral. **B**, The means of the largest areas of infarction were 1.52±0.26 mm2 at POD3 and 0.74±0.08 mm2 at POD14 (p=0.02, two sample t-test). Error bars are defined as SEM.

## vM1 infarction disturbed temporal coding in vS1

To measure the sensory processing of vibrissa inputs, MUA evoked by whisker deflections was recorded at different vS1 depths (Fig. 2A). In L2/3 and deeper L5 (L5b) of sham mice, MUA was precisely time-locked to repetitive whisker stimuli (Fig. 2B). The time-course of the peristimulus time histograms (PSTHs) accurately reported the onset of vS1 inputs. In the acute phase of the vM1 infarction (POD3), MUA increased in the onset response (0–30 ms after onset of stimulus) and the sustained response (30–180 ms after onset of stimulus) (Supplemental Table 1 and Fig. S1), however, MUA in the sustained response was higher than in the onset response. Therefore, the onsets of the next vibrissa inputs were relatively hard to identify from the PSTHs in either L2/3 or L5b (Fig. 2B). These changes in MUA at POD3 lowered the TCI (see Methods for definition of TCI) (Fig. 2C). In the subacute phase of infarction (POD14), the TCI was recovered to the sham level in L2/3 but remained in L5b (Fig. 2C). In fact, among the observed regions, only in L5b the recovery did not occur at POD14 (Supplemental Table 1). These data indicate layer dependency in the M1 infarction effect.

**Figure 2.**
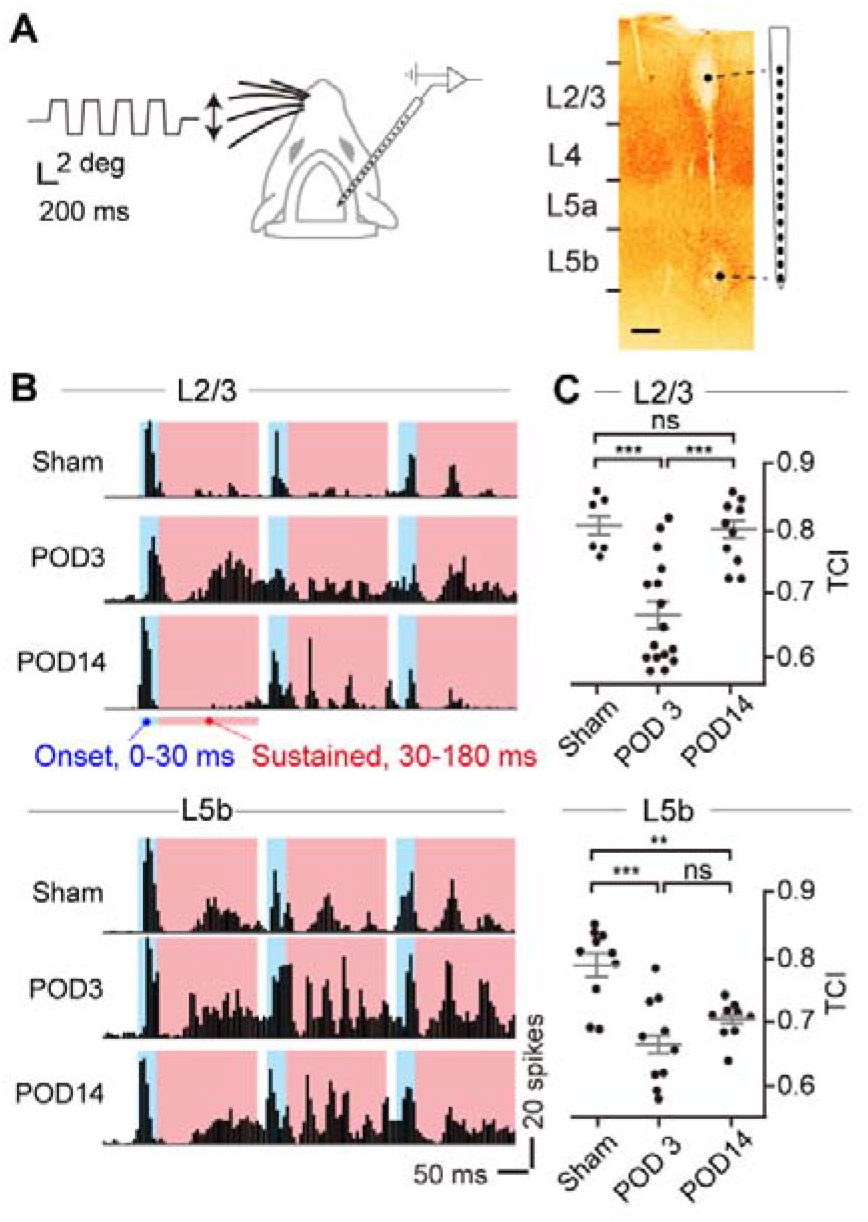
vM1 infarction disturbed temporal coding in vS1 **A**, The whisker stimulation and 16-channel extracellular recording set up. *Left*, Directions of the whisker deflection (double-edged arrow) and a trace of the whisker position. *Right*, Electrolytic lesions at both ends of the recording sites. Scale bar, 100 μm. **B**, Examples of peristimulus time histograms (PSTHs) of multiunit activity (MUA) evoked by the whisker deflections in L2/3 (upper) and L5b (lower) from sham, POD3, and POD14 mice. MUA was classified into the onset (blue area, 0–30 ms after the deflection onset) and sustained (red area, 30–180 ms after the deflection onset) responses. **C**, Temporal coding index (TCI, see Methods) in L2/3 (upper) and L5b (lower) from sham (3 mice), POD3 (5 mice) and POD14 (4 mice). ***, P < 0.001; **, P < 0.01; Tukey’s honestly significant difference test. ns, not significant.

## Laminar patterns of projections from vM1 to vS1

The projection pattern from vM1 to vS1 is not uniform but varies between layers in vS1.^12,13^ To quantify the layer dependency of the projection pattern from the vM1 infarction area, we injected an anterograde tracer, BDA, into vM1 (Fig. 3A) and identified vM1 axonal innervations in vS1. The signal intensity (green line in Fig. 3B) of the anterogradely labelled axons from vM1 was calculated. At the population level, the signal intensity was relatively stronger in L1 and L5b, but weaker in L4 compared to the mean intensity of all layers (N = 3) (Fig. 3C, Fig. S2). As summarised in figure 3D, vM1 axons projected densely to L1 and L5b, where the temporal coding of neural activity did not recover at POD14. This observation suggests that the loss of the synaptic inputs from vM1 to vS1 has an essential role in sensory processing impairment after the vM1 infarction.

**Figure 3.**
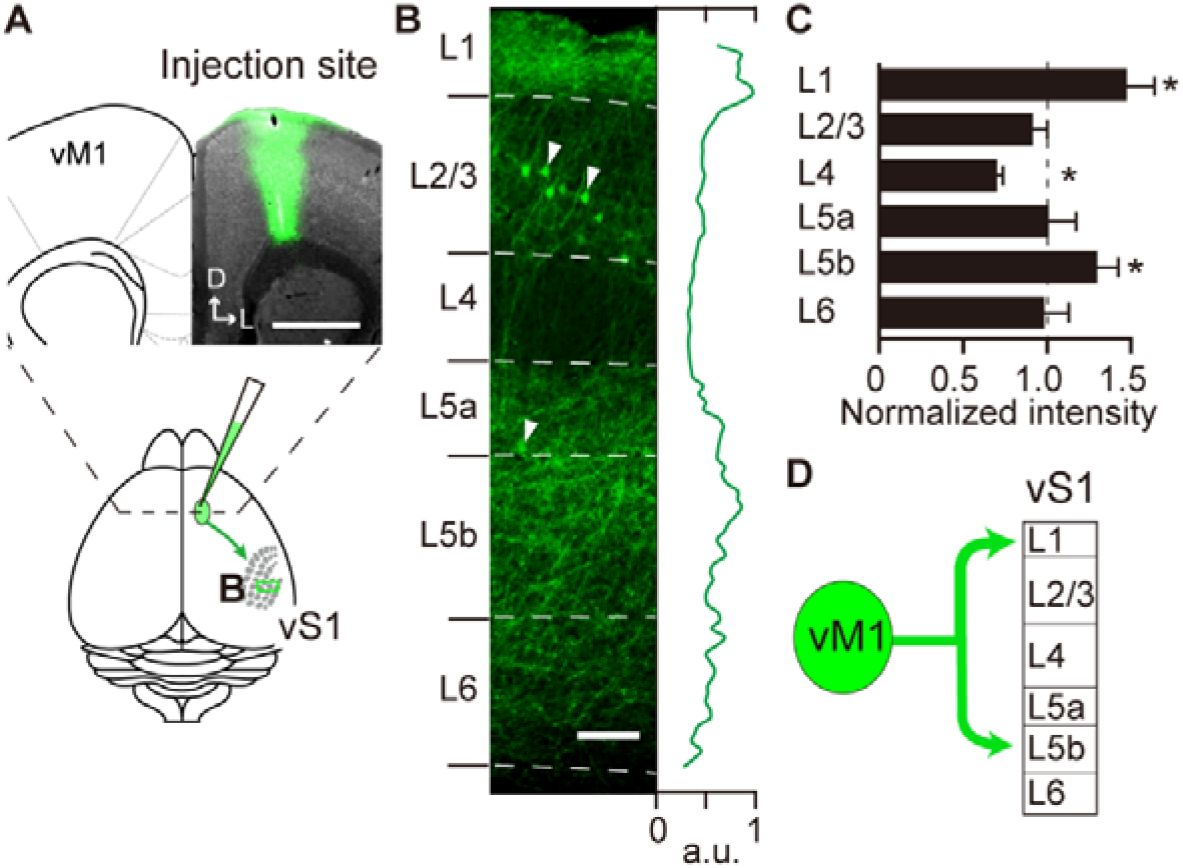
Laminar patterns of vM1 axonal innervations to vS1 **A**, Anterograde tracer injection into vM1. Scale bar, 1 mm. D, dorsal; L, lateral. **B**, Labeled vM1 axons in vS1. The pixel intensity of the axon signals was normalised to the peak value (green line). The signals of retrogradely labelled L2/3 and L5 neurons (arrowheads) were extracted from this measurement. Scale bar, 100 μm. **C**, Laminar distribution of vM1 axons in vS1. Axons intensely innervated L1 and L5b, but sparsely innervated L4 compared to the mean of all layers (*, P < 0.05, one sample t-test; 3 mice). **D**, vM1 axons selectively innervate L1 and L5b of vS1.

## Simulation of vS1 synaptic inputs from vM1

To determine the effect of vM1 on the sustained response in vS1, we first recorded MUA from vM1 and vS1 simultaneously in sham animals (Fig.4A and B). A sensory-evoked vM1 response (Fig. 4A, upper) was observed after the response in vS1 (Fig. 4A, lower) with a constant time delay. Furthermore, each onset and sustained response was longer in vM1 than in vS1. These observations raised the possibility that vM1 receives excitatory inputs from vS1 with a constant delay and integrates them in a constant time period. To elucidate this possibility, we simulated the sensory-evoked vM1 responses from the vS1 responses by using an integrate-and-fire model based on excitatory synaptic connections in the direct pathway from S1 to M1 (see Materials and Methods, Fig. 4B).^13,14^ To maximise the correlation coefficient (*r*) between recorded and simulated vM1 responses, the time window for integration (TW) and the time delay in firing (τ) were estimated to be 30 ms and 6 ms, respectively (Fig. S3). Using these values, the simulated vM1 responses (Fig. 4B, blue line) successfully reproduced the vM1 responses (Fig.4B, gray bars, *r* = 0.68, P < 0.001). Therefore, it is likely that vM1 received sensory inputs directly from vS1, though we cannot rule out the possibility of the trans-thalamic pathway, in which S1 activates M1 through the posterior medial thalamic nucleus (POm).^15^

**Figure 4.**
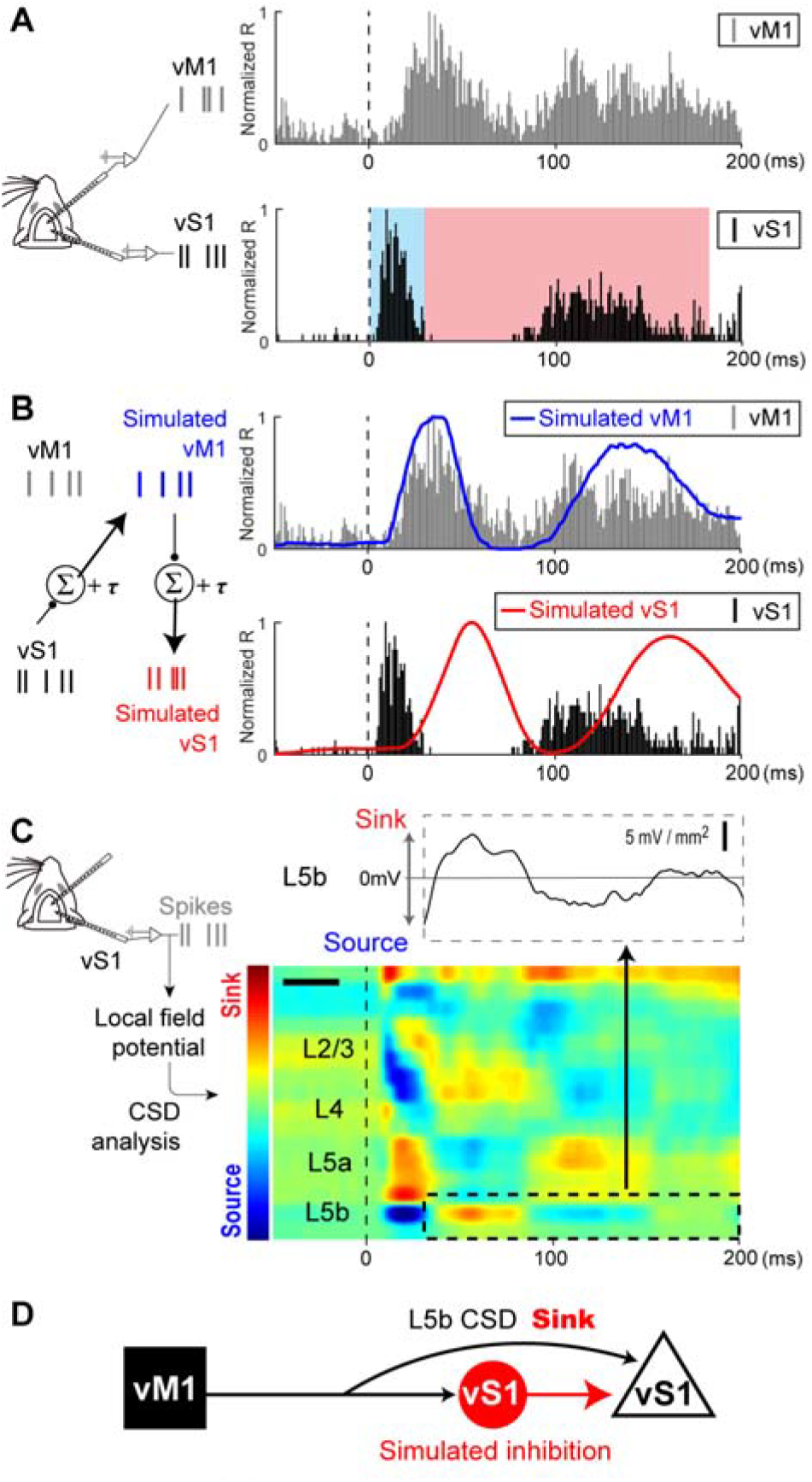
Simulation and CSD analysis indicate inhibition from vM1 to vS1 **A**, Sensory-evoked responses simultaneously recorded from both vS1 and vM1. Recorded vM1 responses (upper, gray bars) and recorded vS1 L5b responses (lower, black bars) to whisker deflections (onset set to zero). **B**, *Left*, A schematic diagram for the simulation of vM1 and vS1 responses using an integrate-and-fire model from recorded vS1 responses. The simulated vM1 responses (blue line) positively correlated with the recorded vM1 responses (gray, the same as in A) (*r* = 0.68, P < 0.001). The simulated S1 responses (red line) negatively correlated with the recorded vS1 responses (black bars, the same as in A) (*r* = −0.52, P < 0.001). **C**, The excitatory synaptic inputs in vS1 were revealed by CSD analysis as current sinks (red in colourmap). Note, L5b of the CSD (dotted area in colourmap and black line in the voltage graph) positively correlated with the simulated S1 responses in **B** (red line, *r* = 0.67, P < 0.001). **D**, A schematic diagram of a network model from the simulation and CSD analysis. The excitatory synaptic input to pyramidal neurons in vS1 (black arrow) was observed as a current sink in the CSD. The input from inhibitory interneurons in vS1 was the simulated vS1 response (red arrow).

We next conducted an additional simulation to predict vS1 responses from the simulated vM1 responses based on the excitatory synaptic connections from vM1 to vS1^13^. For this simulation too, we set TW = 30 ms and τ = 6 ms to the integrate-and-fire model based on excitatory synaptic connections from vM1 to vS1. In contrast to the vM1 simulation, the recorded vS1 responses and the simulated vS1 responses were negatively correlated (Fig. 4B, black bars and red line, respectively, *r* = −0.52, P < 0.001). Although there are both excitatory and inhibitory inputs from vM1 to vS1^16,17^, our results suggest that the net effect from vM1 on vS1 was relatively inhibitory.

To validate the credibility of this simulation, we further performed CSD analysis from the local field potentials recorded from vS1 to confirm if synaptic inputs from vM1 to vS1 exist. From the CSD analysis, we found excitatory inputs patterns to pyramidal neurons, which are considered the most likely generator of the CSD profile (see Methods).^18^ As such, CSD analysis can reveal excitatory synaptic inputs from vM1 as current sinks. The most promising candidate of the excitatory inputs from vM1 was vS1 L5b, where vM1 is innervated densely (Fig. 3C). The time course of the L5b CSD profile (dotted area in Fig. 4C) and that of the simulated S1 responses (red line in Fig. 4B) were positively correlated (*r* = 0.67, P < 0.001). This result indicates the validity of the vS1 simulation from vM1. In sum, it is likely that the net effect of inputs from vM1 is inhibitory via inhibitory interneurons (Fig. 4D).^16,19^

## Inhibition from vM1 after infarction recovered temporal coding in vS1

The results from the simulated vS1 response (Fig. 4) suggest the inhibition from vM1 to vS1 disappeared in the case of the vM1 infarction. To confirm this hypothesis, we tested whether the simulated vS1 response could rescue the temporal coding if virtual inhibition from the simulated vM1 response was applied under the vM1 infarction condition. We first confirmed the loss of the synaptic inputs to the excitatory pyramidal neurons from vM1 by removing the current sink in L5b (Fig. 5A). Then, we applied the simulated inhibition from vM1 to vS1 at POD3 (Fig. 5B). The result was a lowered sustained response and recovered temporal coding. At the population level, the simulated inhibition from vM1 effectively recovered temporal coding to the sham level in L2/3 at POD3 and L5b at POD3 and POD14 (Fig. 5C). These results strongly suggest that the loss of inhibition from vM1 causes deficits in temporal coding in vS1.

**Figure 5.**
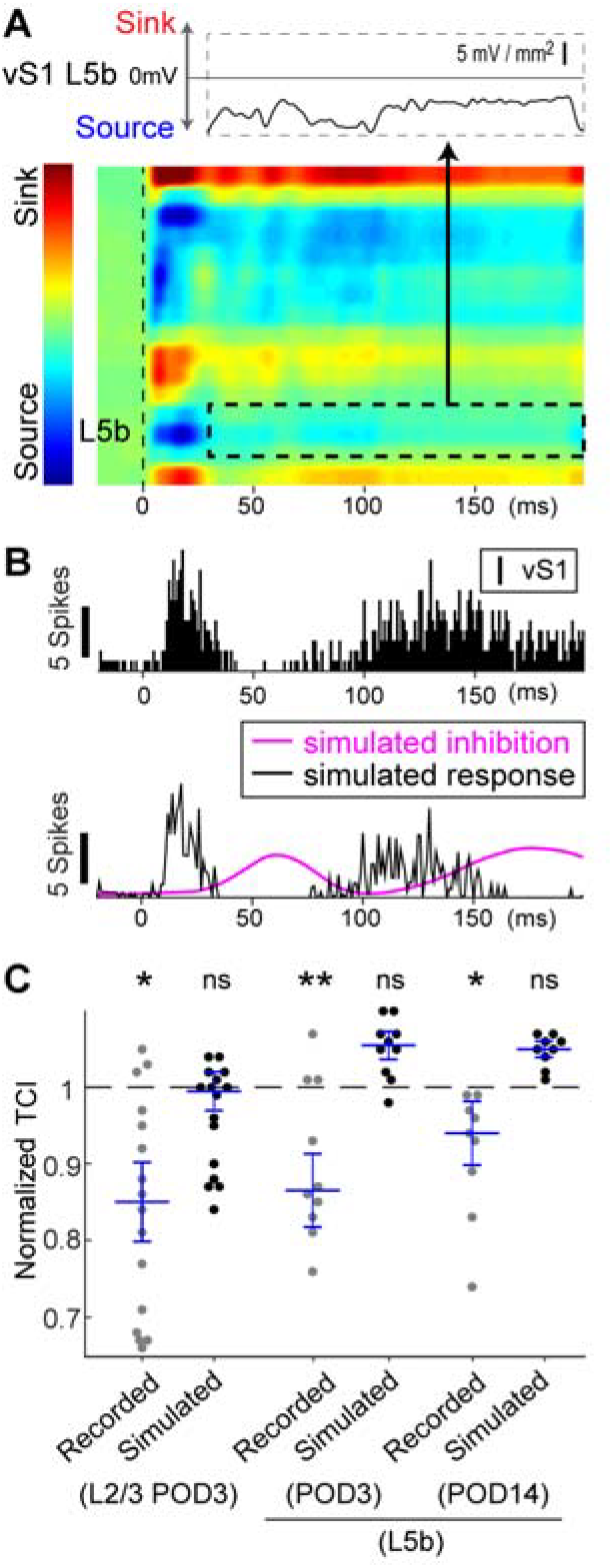
Application of inhibitory input to vM1 infarction can mimic temporal coding in sham animals. **A**, The loss of excitatory inputs to vS1 L5b (dotted area) was visualised by CSD analysis at POD3. **B**, *Upper*, An example of the recorded vS1 L2/3 responses at POD3. *Lower*, Simulated inhibition from vM1 (cyan line) suppressed the sustained responses. **C**, The TCI was recovered to the sham level in simulated vS1 responses: in L2/3, TCI at POD3 of recorded (0.85 ± 0.03) and simulated (1.00 ± 0.03); in L5b, TCI at POD3 of recorded (0.87 ± 0.03) and simulated (1.06 ± 0.01); in L5b, TCI at POD14 of recorded (0.94 ± 0.03) and simulated (1.05 ± 0.01). Each value was normalised to the level of sham mice. *, P < 0.05; **, P < 0.01; Dunnett test compared to the sham level. ns, not significant. For calculating normalised TCI, the response to the first stimulus (from the stimulus onset to 180 ms) was used.

## Discussion

In this study, we showed that the temporal coding of whisker-mediated sensory inputs in vS1 was impaired in the acute phase of the vM1 infarction, but recovered in all layers except L5b in the subacute phase. A tracer study indicated that L5b in vS1 received dense innervation from vM1, while a simulation study and CSD analysis strongly suggested that the vM1 infarction impairs temporal coding in vS1 by the loss of inhibition from vM1 to vS1.

Motor infarction occasionally causes sensory deficits. In the present study, we found that the vM1 infarction increased the sustained response to whisker deflection more significantly than the onset response. This unbalanced increase between the two responses resulted in an impairment in temporal coding (Fig. 2). It has been reported that focal infarction of the neocortex induces disinhibition by widespread alternations in GABA_A_ receptor subtypes at various brain regions^20^, an effect that may underlie the reorganisation of the somatotopy map in S1.^4,5^ Meanwhile, vM1 directly modulates vS1 activity via disynaptic inhibition in normal animals.^16,17^ Thus, these inhibitory mechanisms may be involved in the impairment of temporal cording under the condition of the vM1 infarction. Especially, the disynaptic inhibition mechanism from vM1 may largely engage temporal coding impairment, because the inhibition from simulated vM1 could recover TCI in simulated vS1 responses under the vM1 infarction (Fig. 5). The physiological function of the sustained response in S1 is considered to be responsible for conscious sensory perception ^42,43^. Moreover, it is thought to be a rebound response resulting from recurrent activation of cortical and subcortical circuitry^21^ and controlled by connected areas^22^, such as inputs from vM1.^23,24^. These studies support our result that the vM1 infarction largely influences the sustained responses in vS1.

The recovery of temporal coding after the vM1 infarction was similar to the data of the recent clinical study showing that sensory deficits observed in acute phase of stroke largely recover over time.^25^ However, the recovery process of our data was not uniform among layers. In the subacute phase, TCI in L2/3 was recovered to the sham level, but TCI in L5b was not (Fig. 2). What is responsible for the difference in the recovery process among layers? A substantial amount of studies have shown that neural repair after stroke depends on tissue adjacent to or connected with the infarct lesion.^26^ vS1 receives inputs not only from vM1 but also from the secondary somatosensory cortex (S2) and POm. vM1, S2, and POm have axons that ramify in L1 of vS1 and overlap with the apical dendrites of L2/3 pyramidal neurons.^9,23,27^ Thus, we speculate that the effect of M1 infarction could be easily compensated by direct inputs from S2 or POm. In contrast, vM1 axons reside mostly in deep L5b and L6, whereas S2 axons comparatively project to L5a strongly, and L6 and POm axons are concentrated in L5a^16,28–31^ These observations support the CSD analysis data that showed the reduction of excitatory inputs in L5b after infarction, suggesting a loss of inputs from vM1 after infarction (Fig. 5A). Although the apical dendrite of L5 neurons arbours within L2/3 and remodels its synapses,^32^ the loss of inputs to L5b after M1 infarction might be hard to compensate from other areas. As such, we propose that the layer-specific connectivity affects the recovery process from M1 infarction.

Sensory inputs are necessary for the successful execution and acquisition of skilful voluntary movements.^3^ Therefore, re-establishing sensory processing and sensorimotor interactions in the infarction-damaged motor system appears to be essential for improving motor function. There is evidence indicating that sensory electrical stimulation (SES) improves motor function after infarction.^2^ However, the optimal timing and protocol of SES are still debated. The reason why SES at 10-30 Hz increases corticospinal excitability is unclear.^33^ As shown in Fig. S4B, it takes about 42 ms from stimulation-evoked S1 responses to receive inhibition from M1 (21 ms from S1 to M1 and 21 ms from M1 to S1). This time corresponds to 24 Hz (1 / 0.042 s). Considering the 24Hz from the viewpoint of S1 inhibition from M1, SES at 30 Hz may be close to the threshold that effectively activates the cortico-cortical reciprocal circuit between M1 and S1. On the contrary, SES greater than 30 Hz does not excite M1-S1 circuits effectively, suggesting why 100 Hz SES is less effective than 10-30 Hz SES at exciting corticospinal neurons.^33^

## Methods

### Animals

All surgical procedures and postoperative care were performed following guidelines of the Animal Care and Use Committee of Tokyo Women’s Medical University. The animal experiment permission was approved under the number AE17-127. All experiments were performed in accordance with relevant guidelines and regulations of the Animal Experiment Ethics Committee of Tokyo Women’s Medical University. Every effort was made to minimize the number and suffering of animals used in this study. The animals were housed in a room maintained at 23 ± 1 °C with a 12 h light/dark cycle. Food and water were available *ad libitum*.

### Photothrombotic infarction in mouse vM1

Male C57BL/6 mice (Sankyo Lab. Service Corp., Tokyo, Japan) 8-12 weeks old were operated. Each animal was anaesthetized with an intraperitoneal injection of ketamine (100 mg/kg) and xylazine (16 mg/kg) mixture and held in a stereotaxic apparatus. The skull was exposed and kept wet with saline on the surface to increase the transparency of the skull. Five minutes after the intraperitoneal injection of 1% rose bengal (100 mg/kg; Wako, Tokyo, Japan), green light coupling with an optic fibre (532 nm wavelength, 0.2 mm diameter, 4.5 mW; Thorlabs Inc., Newton, NJ, USA) was applied for 15 min to the right vM1 (1.4 mm anterior to the bregma and 1.1 mm lateral to the midline^12,13^) (Fig. 1).^10^ Subsequently, a head plate was glued onto the skull, and the animal was returned to the home cage. Infarct volumes were calculated using ImageJ by measuring the largest area of the infarct in all coronal sections, which was normalised by the ratio of ipsilesional to contralesional cortical volumes to exclude the shift effect of cortex.

### *In vivo* electrophysiological recording

Electrophysiological recordings were performed at postoperative day 3 (POD3) and POD14, which correspond to the acute and subacute phases of the infarction, respectively (Fig. 1). These times are analogous to the acute (<30 days, corresponding to the inpatient rehabilitation period) and subacute (60-90 days, corresponding to the typical outpatient therapy delivery) phases in human.^34^ Each mouse was anaesthetized with isoflurane (0.8-1.0% during recording) supplemented with an intraperitoneal injection of chlorprothixene hydrochloride (2 mg/kg) for sedation. The respiration rate was monitored and maintained at 100–120 breaths per minute by using a custom-made respiration monitor, 30 Hz USB camera and an acceleration monitor to detect the rib cage motion of the animal. The animals were maintained at 37.0□ rectal temperature by a feedback-controlled heating pad. A silicone probe with 16 recording sites spaced 50 μm apart (A1×16-5mm-50-703; NeuroNexus, Ann Arbor, MI, USA) was inserted into vS1, which corresponded to the C3 or D3 barrel (1.7 mm posterior to the bregma and 3.5 mm lateral to the midline, 35 degrees inclined laterally) until the tip reached a 900 μm depth from the pial surface. Then, local field potentials and multi-unit activities (MUA) were obtained simultaneously from different cortical depths.

### Spike Detection

Data were recorded using a multi-channel acquisition processor (Plexon Inc., Dallas, TX, USA). Local field potentials and multi-unit activity were separated using band-pass filtering at 0.5–300 Hz and 300–6000 Hz, and sampled at 1 kHz and 40 kHz, respectively. At the end of the recording, electrolytic lesions were made at the surface and the deepest recording sites by delivering a small positive current (3 μA, 10 sec) to determine the exact laminar localisation of recording sites.^35^ For spike detection, a threshold was determined by following equation^36^:

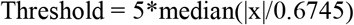

Where x is the bandpass filtered signal. The spike number as multi-unit activity was counted and used for making peri-stimulus time histogram (PSTH). In one animal, both M1 and S1 responses were recorded simultaneously by using two silicone probes (A1×16-5mm-50-703) for checking the validity of the simulation.

### Whisker stimulation

Whisker stimulations were generated using a piezoelectric device controlled by a custom-written MATLAB program (MathWorks, Natick, MA, USA). All whiskers were deflected forward and backward with a 170-ms steady state after each deflection. This routine was cycled four times in one session, and the session was repeated 40 times at 3.6-s intervals. To evaluate the selectivity to deflection onset, the temporal coding index (TCI) was calculated as follows:

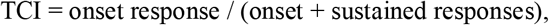

where the onset and sustained responses were the spike numbers during 0 – 30 ms and 30 – 180 ms after the whisker deflection. The responses to every deflection were used to calculate TCI. A value of TCI close to one indicates that the MUA is time-locked to the onset of the deflection.

### Histological identification of laminar positions of recording sites

At the end of the recording, electrolytic lesions at both ends of the recording sites were made by delivering a small positive current (3 μA, 10 sec) to determine the exact laminar localisation of the recording sites. After the recording, the mouse was deeply anaesthetized with sodium pentobarbital (60 mg/kg, intraperitoneally) and transcardially perfused by a fixative solution (4% paraformaldehyde and 0.2% picric acid in 0.1M phosphate buffer). The brain was removed and post-fixed overnight at 4°C. The brain was cut into 40-μm coronal sections using a vibratome (Leica VT1200S; Leica Microsystems, GmbH, Wetzlar, Germany). Sections were incubated overnight at 4°C or 2–3 h at 37°C with 0.05% 3, 3’-diaminobenzidine, 0.03% cytochrome c oxidase, and 4% sucrose in 0.1 M phosphate buffer, mounted on glass slides, and coverslipped using Eukitt (ORSAtec GmbH, Bobingen, Germany). Cortical layers in S1 were identified as follows using cytochrome c oxidase activity: L4 has visible barrel structures; L2/3 has a lower signal than L4; L5a has a lower signal than L4, and L5b has a higher signal than L5a.

### Anterograde labelling from vM1 to vS1

Biotinylated dextran amine (BDA; molecular weight 10,000; 10% in saline; Thermo Fischer Scientific, Waltham, MA, USA) was injected into vM1 (1.4 mm anterior to the bregma and 1.1 mm lateral to the midline^12,13^). After a survival period of seven days, the mouse was deeply anaesthetized with sodium pentobarbital (60 mg/kg, intraperitoneally) and transcardially perfused by a fixative solution (4% paraformaldehyde and 0.2% picric acid in 0.1M phosphate buffer). The brain was cut coronally into 40-μm sections. Sections were incubated overnight with a guinea pig monoclonal antibody against vesicular glutamate transporter type 2 (VGluT2) (1:500; VGluT2-GP-Af810; Frontier Institute Co., ltd., Ishikari, Japan) followed by Alexa Fluor 594-conjugated secondary antibody (for VGluT2; 1:500; Jackson ImmunoResearch, West Grove, PA, USA) and Alexa Fluor 488-conjugated streptavidin (Thermo Fisher Scientific), and subsequently with NeuroTrace 435/455 (1:100; Thermo Fisher Scientific). Cortical layer structures in vS1 were identified using cytoarchitecture, and the VGluT2 staining pattern was similar as previously reported (see Fig. S2).^37^ Three serial sections were used to evaluate the pixel intensity of anterogradely labelled axons along the centerline of the barrel structure of vS1. The pixel intensity in each section was averaged and standardised with maximum values (three mice). These values were further normalised to the mean value of all layers.

### Image acquisition and identification of S1 layer structures

Images were detected by a Zeiss epifluorescence microscopy (Axio Scope.A1, Carl Zeiss, Oberkochen, Germany) equipped with a cooled CCD camera (RS 6.1, Quantum Scientific Imaging, Inc., Poplarville, MS, USA), acquired using a μManager (http://www.micro-manager.org) and ImageJ software (https://imagej.nih.gov/ij), and saved as TIFF files. The contrast and brightness of the images were modified using an Adobe Photoshop software (Adobe Systems Inc., San Jose, CA, USA). Layer structures in S1 were identified using cytoarchitecture and VGluT2 staining pattern as follows similar to the previous report^38^: layer 1 (L1) has few cell bodies; L2/3 has a weaker VGluT2-staining than L4; L4 has a VGluT2-positive barrel structures; L5a has a weaker VGluT2-staining than L4 and L5b, and L6 has smaller cell bodies than L5.

### Simulation analysis of inputs from vM1

To evaluate the role of vM1 in modulating S1 responses,^16,22,24^ we simulated the effect of M1 inputs in two steps. First, we determined the parameters that explain well the responses transferred from S1 to M1 and vice versa. For this step, vM1 responses were simulated from the vS1 responses and compared with the recorded vM1 responses. Second, the effect of vM1 was simulated from the simulated vM1 responses. In both steps, the simulated responses were calculated using the integrate-and-fire model.^14^

The simulated vM1 response (M1simR) was calculated as a summation of vS1 responses (S1R) from the following equation:

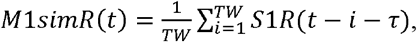

where TW is the time window (ms) for integrating the presynaptic inputs, i.e., summation of spike activity, and r is the time delay to fire (ms) between vM1 and vS1 responses.

The simulated S1 response (S1simR) was calculated from the following equation:

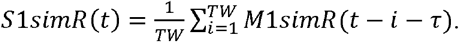

### Current source density analysis

Current source density (CSD) analysis was used to detect the time and the cortical layer of the synaptic inputs, which was observed as the current sink.^18^ At a certain cortical depth (z), the relation between the estimated CSD, *Ĉ*(*z*), and measured potential, Ø(*z*), can be estimated as follows:

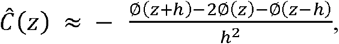

where h is the distance between adjacent recording sites (50 μm in this study). Recording trials were repeated 40 times in each experiment to obtain averaged CSD. We estimated the CSD at the top and bottom electrode contacts by the method of Vaknin et al.^39^ A three-point Hamming filter was used to decrease spatial noise.^40^ Data are represented as a pseudo-colour code from red (sink) to blue (source).

### Statistical Methods

Data are represented as the mean ± standard error (SEM). Statistical analyses were performed using SPSS 25.0 (SPSS Inc., Chicago, IL, USA). The specific tests used are stated alongside all probability values reported.

## Acknowledgements

We thank Dr Goichi Miyoshi for discussions and comments on the manuscript; Dr Hidetsugu Asano for developing a camera-based respiration monitor; and Yumi Tani and Sachie Sekino for technical assistance.

## Author Contributions Statement

A.F., H.O., and M.M. designed the experiments. A.F., H.O., and Y.U. performed and analysed experiments. A.F., H.O., Y.U., and M.M. wrote the manuscript. Y.M. and T.K. supervised all studies. All authors reviewed the manuscript.

## Data Availability

The datasets generated during and/or analysed during the current study are available from the corresponding author on reasonable request.

