## Supplemental material for "Layer-specific sensory processing impairment in the primary somatosensory cortex after motor cortex infarction"

**Supplemental figure**

**Figure S1**

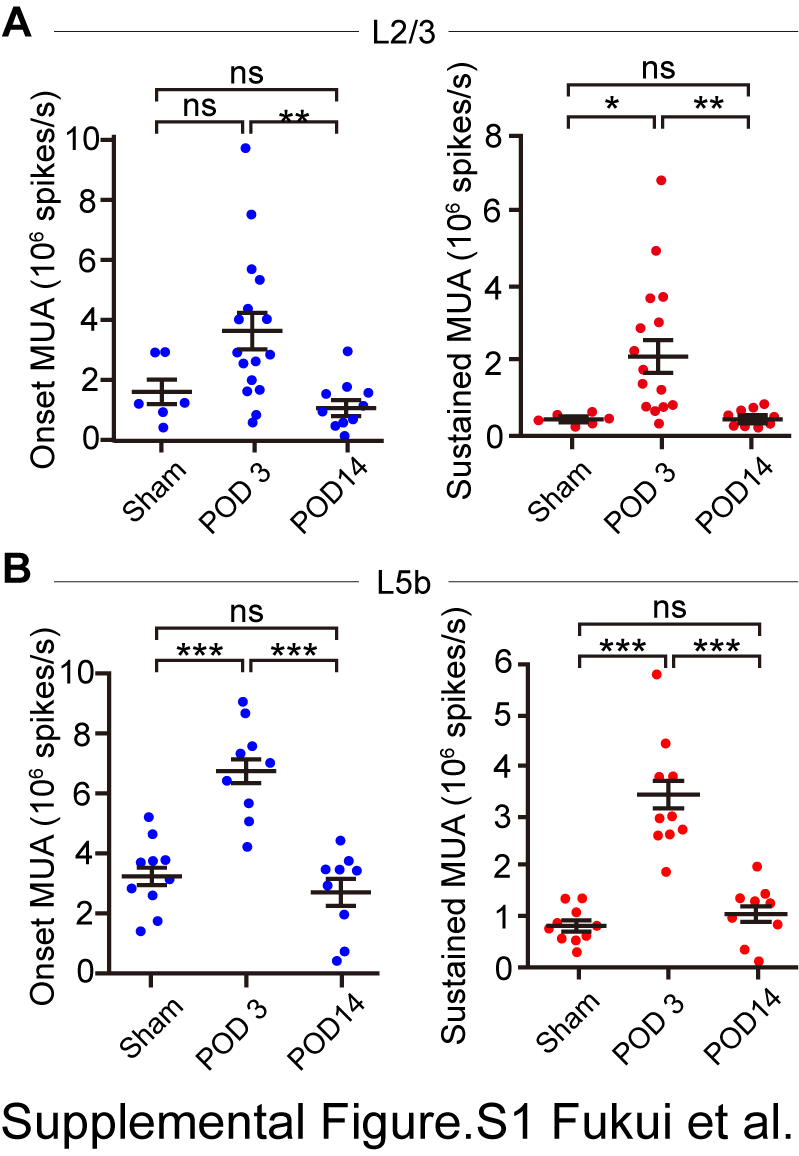

**A**,　MUA evoked by whisker deflection at the onset (0–30 ms) and sustained (30–180 ms) in L2/3 of sham (6 recordings from 3 mice), POD3 (16 recordings from 5 mice) and POD14 (10 recordings from 4 mice). **B**, MUA in L5b of sham (10 recordings from 3 mice), POD3 (10 recordings from 4 mice)　and POD14　(9 recordings from 5 mice). ***, P < 0.001; Tukey-Kramer test. ns, not significant.

**Figure S2**

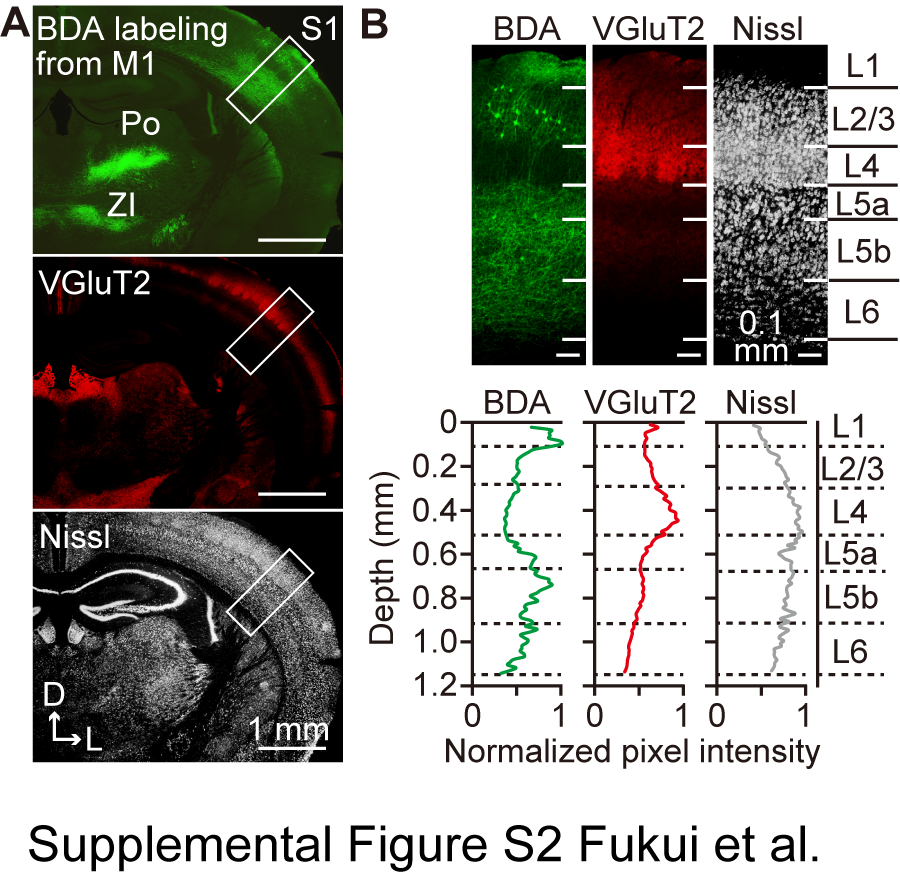

**A**, Anterogradely-labeled axons by BDA (biotinylated dextran amine) injection from vibrissa M1. Vibrissa S1 area and its layer structure were identified by VGluT2 and Nissl staining (B). Note that labelled axons in POm and zona incerta (ZI) indicate exact localisation of tracer injection in M1. **B**, Identification of S1 layers by VGluT2 and Nissl staining. The intensity of fluorescent signals for BDA, VGluT2, and Nissl across layers.

**Figure S3**

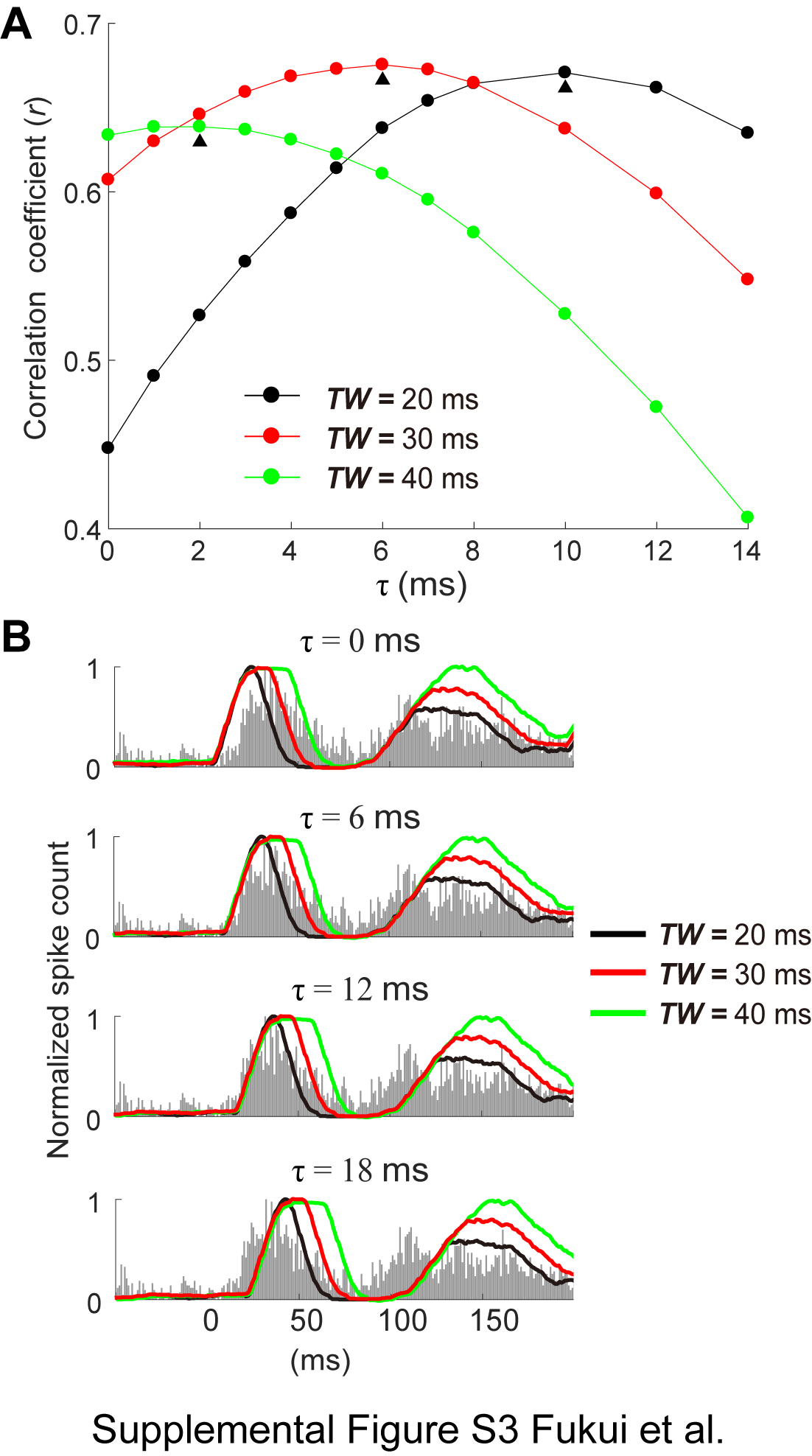

**A**, The values of the correlation coefficient (*r*) between recorded and simulated vM1 responses varied according to the parameters in the integrate-and-fire model. The *r* is highest at the integration time window (*TW*) was 30 ms the delay to fire (τ) was 6 ms. Triangles indicate peak points of the *r* in each *TW*. The *r* was calculated between 0 to 200ms from the stimulus onset. **B**, PSTHs for the simulated M1 responses at various *TW* and τ overlaid on PSTHs for the recorded ones (grey bars, 1 ms/bin).

**Figure S4**

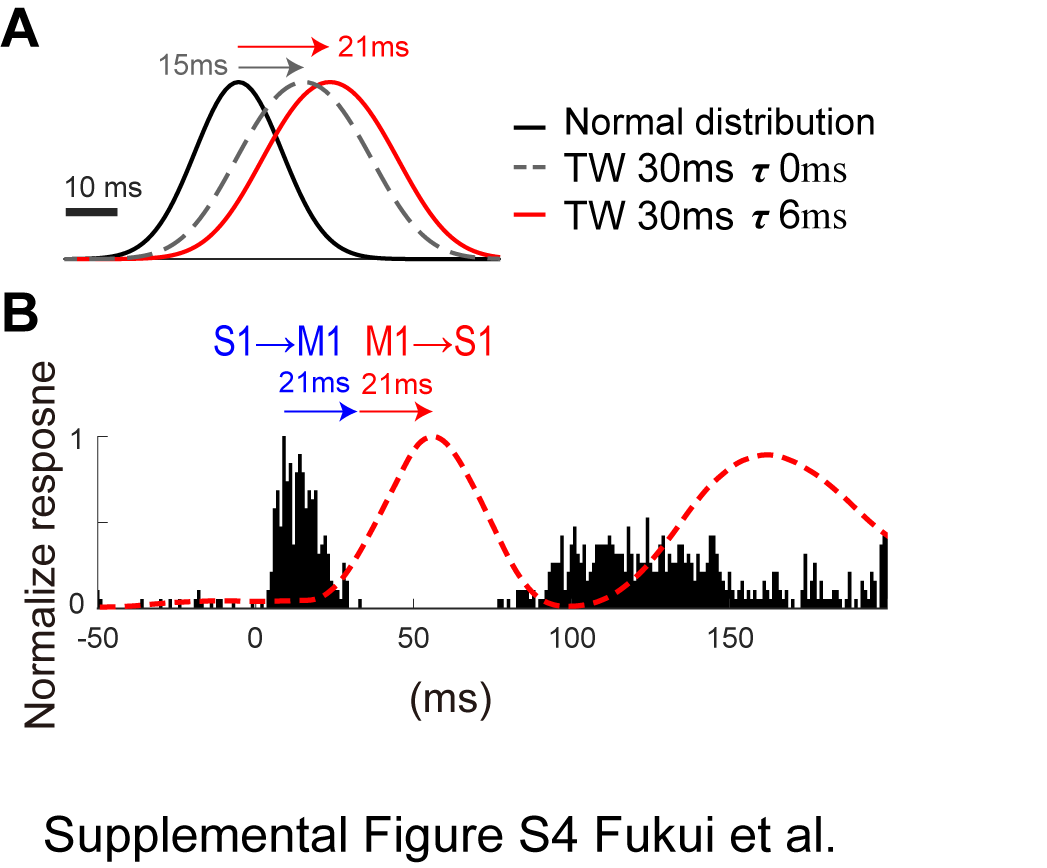

**A**, Simulated S1 responses was delayed 21 ms from the previous stage, i.e., M1 responses when *TW* and *τ* were fixed to 30 ms and 6 ms, respectively. **B**, The simulated inhibitory effect from M1 (red dotted line) was delayed 42 ms (21+21ms) from recorded S1 responses (black PSTH).

**Supplement table**

Table 1. Multiunit activity (MUA) recorded from different layers of mouse vS1

|  | MUA | sham | poststroke | poststroke | Statistical comparisons | | |
| --- | --- | --- | --- | --- | --- | --- | --- |
|  | (10^6^spikes/s) | (a) | (b) POD3 | (c) POD14 | a vs. b | b vs. c | a vs. c |
| L2/3 | Onset MUA | 1.71 ± 0.43  (n = 6) | 3.76 ± 0.62  (n = 16) | 1.29 ± 0.26  (n = 10) | n.s.  (P = 0.078) | b > c  (P = 0.008) | n.s.  (P = 0.900) |
|  | Sustained MUA | 0.34 ± 0.06  (n = 6) | 2.07 ± 0.43  (n = 16) | 0.31 ± 0.10  (n = 10) | a < b  (P = 0.018) | b > c  (P = 0.004) | n.s.  (P = 0.999) |
|  | Onset/total ratio | 0.82 ± 0.02  (n = 6) | 0.67 ± 0.02  (n = 16) | 0.80 ± 0.02  (n = 10) | a > b  (P < 0.001) | b < c  (P < 0.001) | n.s.  (P = 0.925) |
| L4 | Onset MUA | 3.95 ± 0.48  (n = 8) | 6.99 ± 0.53  (n = 17) | 4.00 ± 0.46  (n = 12) | a < b  (P = 0.002) | b > c  (P < 0.001) | n.s.  (P = 0.998) |
|  | Sustained MUA | 0.99 ± 0.15  (n = 8) | 4.15 ± 0.53  (n = 17) | 1.53 ± 0.12  (n = 12) | a < b  (P < 0.001) | b > c  (P < 0.001) | n.s.  (P = 0.717) |
|  | Onset/total ratio | 0.80 ± 0.01  (n = 8) | 0.65 ± 0.02  (n = 17) | 0.71 ± 0.02  (n = 12) | a > b  (P < 0.001) | b < c  (P=0.015) | a > c  (P = 0.006) |
| L5a | Onset MUA | 3.61 ± 0.50  (n = 6) | 5.66 ± 0.44  (n = 11) | 3.19 ± 0.49  (n = 8) | a < b  (P = 0.022) | b > c  (P = 0.003) | n.s.  (P = 0.837) |
|  | Sustained MUA | 1.34 ± 0.24  (n = 6) | 3.69 ± 0.39  (n = 11) | 1.46 ± 0.23  (n = 8) | a < b  (P < 0.001) | b > c  (P < 0.001) | n.s.  (P = 0.969) |
|  | Onset/total ratio | 0.74 ± 0.02  (n = 6) | 0.61 ± 0.02  (n = 11) | 0.68 ± 0.01  (n = 8) | a > b  (P < 0.001) | b < c  (P = 0.011) | n.s.  (P = 0.106) |
| L5b | Onset MUA | 3.30 ± 0.38  (n = 10) | 6.69± 0.49  (n = 10) | 2.73 ± 0.47  (n = 9) | a < b  (P < 0.001) | b > c  (P < 0.001) | n.s.  (P = 0.662) |
|  | Sustained MUA | 0.86 ± 0.11  (n = 10) | 3.38 ± 0.36  (n = 10) | 1.11 ± 0.19  (n = 9) | a < b  (p < 0.001) | b > c  (p < 0.001) | n.s.  (P = 0.767) |
|  | Onset/total ratio | 0.79 ± 0.02  (n = 10) | 0.67 ± 0.02  (n = 10) | 0.70 ± 0.01  (n = 9) | a >b  (P < 0.001) | n.s.  (P = 0.242) | a > c  (P = 0.007) |

Data are mean ± SEM.

Onset MUA, MUA of <30 ms from whisker stimulation; sustained MUA, MUA of 30–180 ms from stimulation; total, summation of onset and sustained MUA. n, number of recorded channels from 3, 5 and four mice for sham, postoperation day 3 (POD3), and POD14, respectively. P value, Tukey–Kramer test. n.s., not significant.
